# Synthesis and Characterisation of Aqueous Haemoglobin-based Microcapsules Coated by Genipin-Cross-Linked Albumin

**DOI:** 10.1101/818278

**Authors:** Kai Melvin Schakowski, Jürgen Linders, Katja Bettina Ferenz, Michael Kirsch

## Abstract

Bovine serum albumin (BSA)-coated haemoglobin (Hb)-microcapsules prepared by co-precipitation of Hb and MnCO_3_ may present an alternative type of artificial blood substitute. Prepared microcapsules were analysed by Scanning electron microscopy (SEM) and Respirometry, cytotoxicity was evaluated by addition of microcapsules to murine fibroblast-derived cell line L929 (American Type Culture Collection, NCTC clone 929 of strain L). The capsules come along with a mean diameter of approximately 0.6 μm and a mean volume of 1.13 ∙ 10^−19^ L, thus an average human red blood cell with a volume of 9 ∙ 10^−14^ L is about 800,000 times bigger. Hb-microcapsules are fully regenerable by ascorbic acid and maintain oxygen affinity because oxygen is able to pass the BSA wall of the capsules and thereby binding to the ferrous iron of the haemoglobin entity. Therefore, these microcapsules present a suitable type of potential artificial haemoglobin-based oxygen carrier (HbOC).

## 1. Introduction

Every day more than 200,000 units of erythrocyte concentrate are needed for transfusion after fatal accidents or during surgery involving the loss of huge amounts of blood. Despite the annual need for more than 85 million units of packed red blood cells (RBCs) worldwide (e.g. about four million units in Germany and 12-16 million units in the USA), a constant decline in willingness to donate blood therefore brings up severe difficulties to handle (Riedel *et al.* 2000, Diez-Silva *et al.* 2010, Sharma *et al.* 2011, Müller *et al.* 2015, Remy and Spinella 2016, García-Roa *et al.* 2017, Henseler 2019). Besides the well-known limits of compatibility among the different blood groups, logistical factors achieve increased importance (Chen *et al.* 2009). Since hospitals must ensure adequate supply of RBCs (and of other blood components) for transfusion and because the number of blood donations is retrograde in highly developed countries, medical institutions must import RBCs and this can evoke a bottle-neck of the supply when the travel time of the units determines the availability.

Shortage on erythrocytes always coincides with shortage on long-term oxygen supply, as a sufficient quota of haemoglobin must be assured in order to secure adequate sustenance. Due to scarcity on RBCs, alternative ways of sustaining blood oxygen levels must be deduced. Within the last years artificial blood substitutes have gained more and more attention because of the above mentioned decrease of available RBCs (Ruchalla 2013, Njoku *et al.* 2015, Chung *et al.* 2016, Ellingson *et al.* 2017, Taguchi *et al.* 2017, Ferenz and Steinbicker 2019). Since the application of a pure solution of haemoglobin is impractical due to the enormous nephrotoxicity and short circulation time (Chang 2006), several different attempts have been made to create biocompatible artificial oxygen carriers, based on either perfluorocarbons or haemoglobin that needs to be encapsulated in protective shells in any case (Bauer *et al.* 2010, von Storp *et al.* 2012, Ferenz *et al.* 2013, Sakai *et al.* 2013, Stephan *et al.* 2014, Wrobeln, Laudien, *et al.* 2017, Wrobeln, Schlüter, *et al.* 2017).

The most commonly used assay for cross-linkage of albumin is the method of glutaric dialdehyde cross-linkage (Ling 1961, Mamedova *et al.* 2002, Choi *et al.* 2006, Bychkova *et al.* 2013). This method bears the disadvantage that excess toxic glutaric dialdehyde must be removed from the reaction mixture by using LiAlH_4_ or NaBH_4_, both highly inflammable chemicals in combination with aqueous environments.

In the present study, we focused on a technique that makes use of the high loading capacity of inorganic carbonates. By co-precipitation of haemoglobin and MnCO_3_ followed by adsorption of cross-linked albumin on the surface of the MnCO_3_, haemoglobin is captured within a shell of albumin, that was cross-linked with genipin or derivates of benzoic acid. Dissolution of the MnCO_3_ template finally leads to porous particles of cross-linked albumin with encapsulated haemoglobin. Although the technique of cross-linking proteins by genipin is an established one (Butler *et al.* 2003, Yoo *et al.* 2011), the preparation of genipin linked albumin shells as part of artificial oxygen carriers is a novel procedure.

## 2. Experimental

### Materials

Ascorbic acid, 4-Bromomethyl-3-nitrobenzoic acid (BNBA), Bovine serum albumin fraction V (BSA), Dimethyl sulfoxide anhydrous (DMSO), 4-(4,6-Dimethoxy-1,3,5-triazin-2-yl)-4-methylmorpholinium chloride (DMT-MM), Ethylenediaminetetraacetic acid disodium salt dihydrate (Na_2_EDTA), Haemoglobin from bovine blood (Hb), Manganese(II) chloride tetrahydrate, Minimum Essential Medium Eagle (MEM), sodium dithionite (SDT) and Triton X-100 were purchased from Sigma-Aldrich (Darmstadt, Germany); Sodium bicarbonate, Sodium carbonate anhydrous, Sodium hydroxide, Potassium cyanide, Potassium ferricyanide and Monopotassium phosphate were purchased from Merck (Darmstadt, Germany); Boric acid, Glucose, *L*-Glutamine, Sodium chloride, Sodium dodecyl sulphate ultra-pure (SDS), Disodium phosphate dehydrate and Potassium chloride were purchased from Carl Roth (Karlsruhe, Germany); Ringer’s solution was purchased from Fresenius Kabi (Bad Homburg, Germany); Gibco Fetal bovine serum (FBS), Gibco Pen Strep and Gibco 0.05% Trypsin-EDTA were purchased from Thermo Fisher Scientific (Schwerte, Germany).

### Preparation of Hb-Microcapsules

Hb-microcapsules were prepared by modifying methods that were originally developed by XIONG *et al.* (Xiong *et al.* 2012, 2013, Li *et al.* 2017). Since lately cross-linkage of proteins by genipin has gained attention (Butler *et al.* 2003, Yoo *et al.* 2011, Shahgholian *et al.* 2017), the chemical cross-linkers BNBA and DMT-MM used by XIONG *et al.* were replaced by genipin, a natural substance derived from the iridoid glycoside geniposide. Although genipin does not serve as a cross-linker as naturally purposed, it’s chemical properties can be used to link amino-groups (Djerassi *et al.* 1960). The mechanism of linking amino-groups by genipin has first been described by BUTLER *et al.* in 2003 (s. Figure 1) (Butler *et al.* 2003).

**Figure 1.**
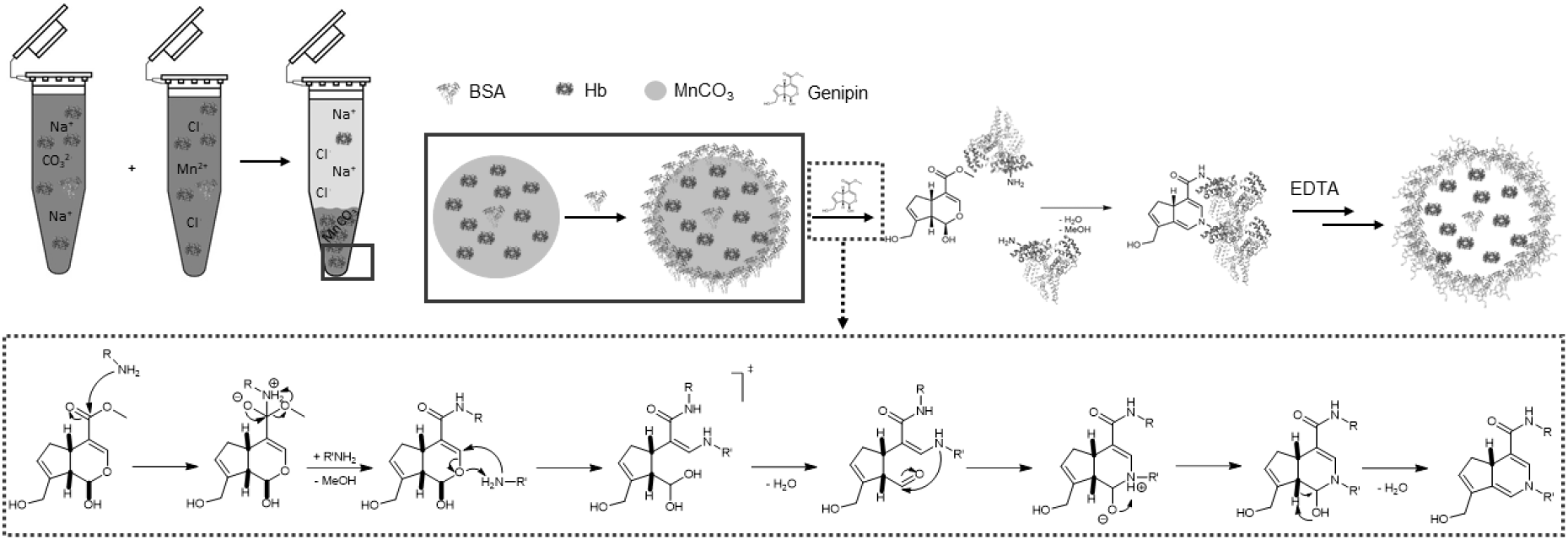
After co-precipitation of MnCO3 and Hb, BSA forms an additional layer around the MnCO3/Hb-particle. BSA within this layer is then cross-linked by addition of genipin and later the MnCO3 in the core of the particle is dissolved by EDTA, resulting in a stable shell of BSA around the Hb-core. The mechanism of cross-linkage by genipin has been described by Butler et al. in 2003. R represents the amino-sidechains of BSA.

A volume of 15 mL of aqueous solutions containing either 0.25 mM Na_2_CO_3_ or MnCl_2_ as well as 10 mg ∙ mL^−1^ Hb and 1 mg ∙ mL^−1^ BSA were mixed under continuous stirring in a 100 mL beaker. After stirring for two more minutes, 150 mg BSA were added slowly under continuous stirring for five more minutes. The resulting suspension of BSA-coated MnCO_3_ particles with entrapped Hb was centrifuged (1000 g, 1 min.). The residue was washed three times with 60 mL of Ringer’s solution each, while the supernatants were collected separately and photometrically checked for their concentrations of remaining Hb. After washing, the residue of particles was suspended in either 60 mL of DMSO containing 37.5 mg BNBA and 45 mg DMT-MM or 60 mL of an aqueous solution of genipin differing from 0.188 – 4 mM genipin. The re-suspended particles were shaken for 24 hours at room temperature. The DMSO-treated fraction was then washed three times with 60 mL DMSO each. From now on both fractions were treated equally. They were washed three times with 60 mL of Ringer’s solution each and then re-suspended in 100 mL of 0.2 M Na_2_EDTA-solution at pH 7.4. The suspension was shaken by hand until all visible solid had completely dissolved. If necessary, samples were sonicated in order to complete the breakdown of the solid. Dissolution of the MnCO_3_ by dissociated Na_2_EDTA resulted in constructs that trapped Hb into the inside of the cross-linked BSA-shell, thus EDTA chelating Mn^2+^ to form a soluble MnEDTA complex. The fractions were centrifuged (1000 g, 1 min.) and the residue was washed two times with 80 mL EDTA-solution, then three times with 60 mL of Ringer’s solution before being re-suspended in 5 mL of Ringer’s solution each and stored at 4 °C under exclusion of light.

### Characterisation of Hb-Microcapsules

#### Photometry

Concentrations of Hb stock solutions were determined by the cyanmethaemoglobin-method (Drabkin and Austin 1932, van Kampen and Zijlstra 1961) as well as the SDS-Hb-method (Oshiro *et al.* 1982, Karsan *et al.* 1993, Kalyan Chakravarthy *et al.* 2012) on a UV-VIS Specord S 600 spectrophotometer (Analytik Jena, Jena, Germany). Concentrations of Hb in the supernatants were determined by the SDS-Hb-method sole, due to residue manganese interfering with the cyanide of the transforming reagent.

For the cyanmethaemoglobin-method, 1 mL of transforming reagent containing 330 mg potassium ferricyanide, 400 mg sodium hydroxide, 150 mg potassium cyanide and 125 mg boric acid per litre was added to 40 μL of aqueous Hb-solutions of different concentrations. The mixtures were incubated for 10 minutes at room temperature and measured against a mixture of 40 μL Millipore water and 1 mL transforming reagent at λ = 546 nm.

For the SDS-Hb-method, 450 μL of Millipore water were added to 50 μL of aqueous Hb-solutions of different concentrations and stirred with 500 μL of 0.06% aqueous SDS-solution. After incubation for 15 minutes at room temperature, the mixtures were measured against 500 μL of Millipore water and 500 μL of SDS-solution at λ = 540 nm.

Since both, cyanmethaemoglobin and SDS-haemoglobin strictly obey the law of LAMBERT and BEER (Lambert 1760), extinction coefficients could be determined by linear regression (n = 5), as depicted in Figure 2. All standard deviations are well below 10^−3^ and hence not shown.

**Figure 2.**
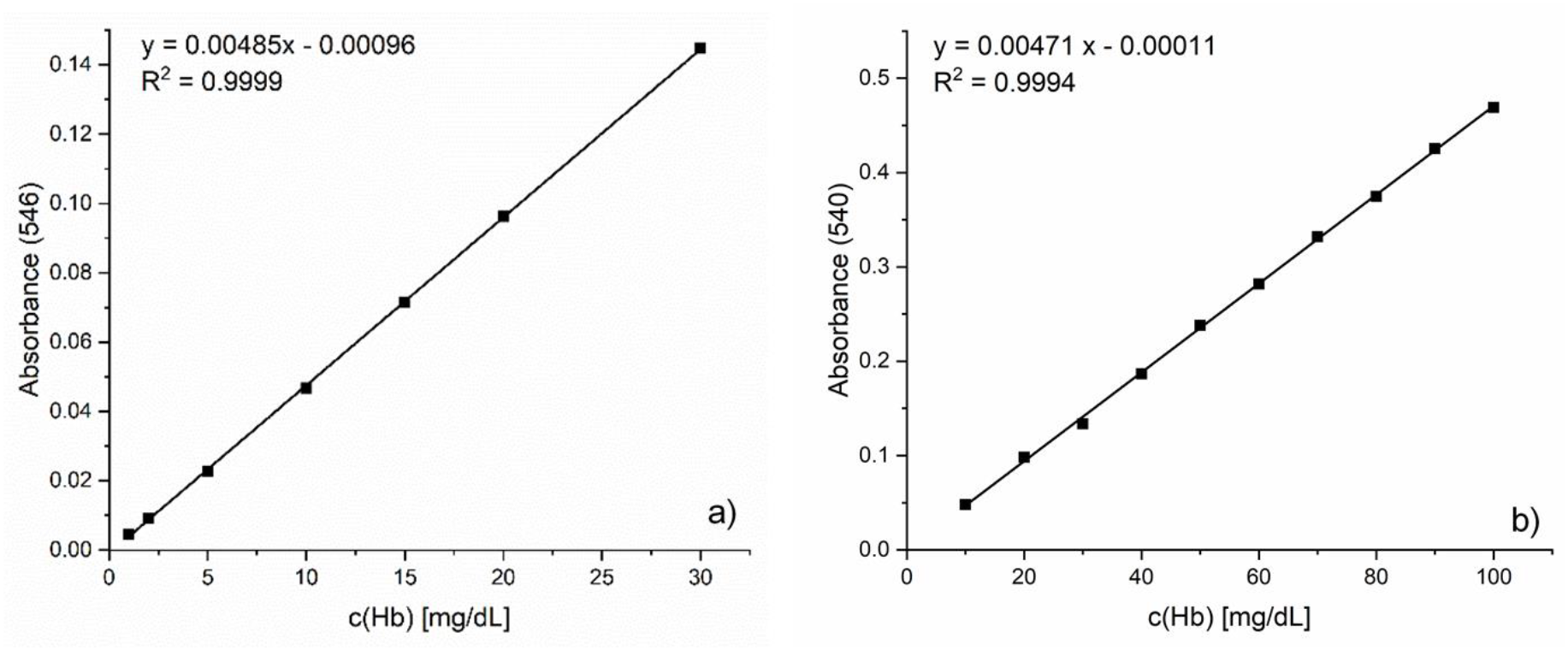
Determination of extinction coefficients for cyanmethaemoglobin(CN)-(a) and SDS-Hb-method (b) by linear regression (n=5 each); Different concentrations of aqueous Hb-solutions were added to transforming reagent (CN) or SDS-solution, respectively. After incubation, absorption of the solutions was measured at 546 nm (CN) or 540 nm (SDS) and plotted against the concentrations.

The entrapment efficiency (EE) was calculated by determining the concentration of Hb in the stock solution ([Hb]_0_) and in the supernatants after co-precipitation with MnCO_3_ ([Hb]_s_) and each washing step. For there was hardly any Hb detectable in the supernatants of each washing step, the EE was calculated according to EE = (1 – [Hb]_s_/[Hb]_0_) ∙ 100%.

#### Respirometry

Respirometry was performed on an Oroboros Oxygraph-2k (Oroboros Instruments, Innsbruck, Austria). In order to determine the oxygen binding capacity of the prepared microcapsules, 1 mL of the suspension of Hb-microcapsules or freshly prepared free Hb solution (same Hb-concentration as in Hb-microcapsules‘ suspension, control) was mixed with 1 mg of sodium dithionite (SDT). The chambers of the Oxygraph were filled with 50 mM sodium phosphate buffer (pH 7.4) and equilibrated with air. After equilibrium of the chambers was reached, 50 μL of the prepared microcapsules’ suspension or free Hb solution was added and the decrease in oxygen concentration within the chamber was measured. The concentration of Hb for each control was calculated individually from the encapsulation efficiency of each sample.

For displaying the regenerative capacity, volumes of a concentrated suspension of Hb-microcapsules and a solution containing 500 μg ∙ mL^−1^ potassium ferricyanide were mixed in ratio 1:2 and incubated over night at 4 °C. The chambers of the Oxygraph were filled with 50 mM sodium phosphate buffer (pH 7.4) and equilibrated with air. After equilibrium of the chambers was reached, 50 μL of the before prepared solution was added. Then 150 μL of a freshly prepared 10 mg ∙ mL^−1^ solution of ascorbic acid was added and decrease of oxygen concentration within the chambers was measured.

#### Dynamic light-scattering

Dynamic light-scattering (DLS) was performed on a Nano-Flex apparatus (Particle Metrix, Meerbusch, Germany) by using Microtrac FLEX 11.0.05 software. 500 μL of the microcapsules’ suspension was transferred into a 2 mL micro reaction vessel (Eppendorf, Hamburg, Germany) and measured three times for 60 seconds against Ringer’s solution for set zero.

#### Scanning Electron Microscopy

Scanning Electron Microscopy (SEM) was performed on a FIB-SEM 540 Crossbeam (Carl Zeiss, Oberkochen, Germany) in the Electron Microscopy Unit of the Imaging Center Essen (University Hospital Essen, Essen, Germany). Samples were prepared by dehydration in an ascending five-step ethanol series from 30-96% followed by immersion in 100% pure dried ethanol three times, for 10 minutes each. Samples were dried by critical point drying with CO_2_ and sputter coated with Pt/Pd. Every image taken was provided with the exact data of capturing.

##### Native Polyacrylamide Gel Electrophoresis

Gradient Polyacrylamide Gel Electrophoresis (PAGE) was used to separate oligomers of BSA from monomer BSA. A gradient from 4% - 16% was used. Proteins were stained by a solution of 2.5 g Coomassie brilliant blue R 250 in a mixture of 500 mL methanol, 100 mL acetic acid completed to 1000 mL using Millipore water. For destaining the same solution without Coomassie was used.

##### Cell Culture and Measurements of Lactate-Deydrogenase (LDH)-Activities

Measurements on LDH-activity and other commonly measured clinical parameters were performed on a Respons®920 (DiaSys Diagnostic Systems GmbH, Holzheim, Germany).

The standard cytotoxicity model of murine fibroblast-derived cell line L929 (American Type Culture Collection, NCTC clone 929 of strain L) was used for the experiments. Cells were cultured in minimum essential medium (MEM) Eagle supplemented with 25 mM sodium bicarbonate, 10% FBS, 2 mM L-glutamine, and 100 units/mL penicillin G and 0.1 mg/mL streptomycin in 75 cm^2^ plastic flasks at 37 °C in a humidified atmosphere of 95% air and 5% CO_2_. Subcultures were obtained by trypsinisation (0.05% trypsin-EDTA) (Lomonosova *et al.* 1998). For experimental use, cells were transferred to 6-well plates.

For toxicity experiments of prepared microcapsules counting of the cells right before beginning of the experiments showed an average number of about 1,000,000 cells per well. The medium was removed and cells were washed with 2.0 mL of PBS buffer before 2.0 mL of the solutions to test were added. All test-solutions contained Ringer’s solution + 4.5 g/L glucose and a) 3 mg/ml BSA (g-BSA) or b) 2.5 – 5 vol% of microcapsules as HbOCs or c) BSA and free haemoglobin in different concentrations (Hb) or d) no further ingredients (blank), respectively.

Wells were incubated at described conditions for 60 – 300 minutes before 1 mL of the supernatant was removed for LDH measurement (LDH_t_, t∈{60, 120, 180, 240, 300}). At the end of the experiment, cells were lysed for 15 min at 37 °C by the addition of 200 μL of 25% Triton X-100 followed by shaking before 1 mL of the remaining solution was removed for LDH measurement to obtain the maximum possible amount of LDH that could have been released (LDH_max_). Total lysis of cells was confirmed by visual inspection under an optical microscope.

All samples were centrifuged shortly to remove larger cell debris and HbOCs before performing photometric measurements on LDH-activity using the automatic LDH protocol of the Respons®920 according to the manufacturer’s instructions (Instructions for use (respons 920) 2019).

For statistical analysis, t-tests were performed assuming two-tailed distributions and unequal variances.

Relative cell damage (d_r_%) has been calculated according to d_r_% = (LDH_t_/LDH_max_) ∙ 100%.

Tests on interference between 5 vol% HbOCs and commonly measured clinical parameters have been performed by diluting standard serums obtained from DiaSys Diagnostic Systems GmbH (Holzheim, Germany) with different quantities of microcapsules solutions or Ringer’s solution, respectively. All samples were centrifuged shortly to remove HbOCs before performing measurements on activities of alanin transaminase (ALT), aspartate transaminase (AST), creatine kinase (CK) and LDH as well as concentrations of creatinine (Crea) and Urea.

##### Quantitative nuclear magnetic resonance measurements

All ^1^H-Nuclear Magnetic Resonance (NMR) diffusion experiments were run on a 500 MHz Bruker Avance NEO II spectrometer with a Bruker DIFF BBI probe head at 298 K (Bruker BioSpin, Rheinstetten, Germany). The free induction decays resulting from the addition of a set of 64 scans (90°-pulse; P1 = 9.57 µs) were Fourier transformed and analysed.

A solution of 20% Triton X-100 in D_2_O was diluted 1:1 with pure D_2_O or a suspension of microcapsules (5 vol%) in D_2_O, respectively, and incubated for five hours at 37 °C. Both samples were centrifuged shortly to remove microcapsules and equal volumes of supernatant were taken for quantitative NMR measurement (qNMR). 7.5 μL of DMSO were added to both samples as internal standard.

## Results and Discussion

Genipin-induced polymerisation of BSA was observable by the change of colour of a shaken solution containing genipin and BSA. The solution of 5% BSA and 0.17% genipin equals a molar ratio of 1:10. At the beginning the solution presented a pale-yellow colour attributable to BSA. After 16 hours of shaking at room temperature the colour had turned into a dark green, the combination of residue BSA monomer and the blue colour of oligomerised BSA described in literature (Touyama *et al.* 1994, Butler *et al.* 2003, Somers *et al.* 2008, Yoo *et al.* 2011). The change in colour was traceable by the appearance of a new signal in the spectrum of the BSA-genipin solution at 599 nm after the applied reaction period, as seen in Figure 3. The pure solution of BSA showed its highest absorbance in the area of blue and ultraviolet light, while the solution of pure genipin displayed hardly any absorbance in the spectrum of visible light. After merging equal volumes of the two solutions, the resulting spectrum presented the absorbance of a diluted solution of BSA. Cross-linking of BSA by genipin was additionally proven by comparing native PAGE traces of pure BSA and genipin-treated BSA to BSA cross-linked by glutaraldehyde, which is widely known to cross-link proteins (Xiong *et al.* 2012, Shahgholian *et al.* 2017). Cross-linking by genipin monitored by native PAGE resulted in a trace virtually identical to that of linking by glutaraldehyde (s. Figure 3).

**Figure 3.**
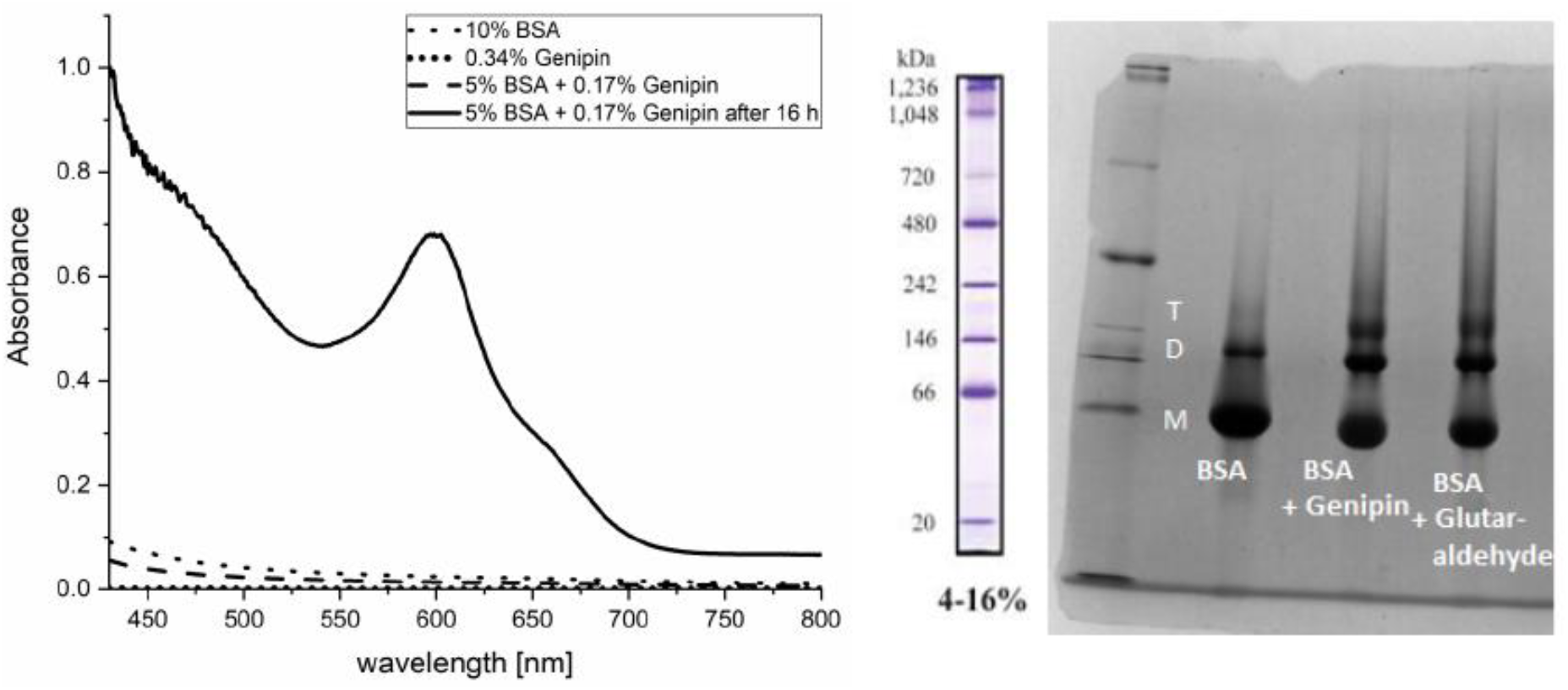
Verification of cross-linking of BSA by genipin. Photometrical spectra of solutions BSA and/or genipin; standardized to a maximum absorbance of 1 (left) and native PAGE of pure BSA, genipin cross-linked BSA and BSA linked by glutaraldehyde as reference (right). M represents monomer BSA, while D represents dimer and T trimer BSA.

The formation of Hb-microcapsules was started by co-precipitation of MnCO_3_ together with Hb, induced by mixing equal volumes of solutions containing either Hb and MnCl_2_ or Hb and Na_2_CO_3_, thus forming MnCO_3_ exceeding its solubility product with a total solubility of 0.065 g ∙ L^−1^ (*GHS Material Safety Data Sheet for MnCl_2_ tetrahydrate* n.d., *GHS Material Safety Data Sheet for MnCO3* n.d., Johnson 1982, Joint Research Centre (European Commission) 2000). Hardly soluble MnCO_3_ particles rapidly formed around Hb lead to entrapment of Hb within these precipitating particles. The subsequent addition of BSA then created a shell around MnCO_3_ particles by adsorbing onto the surface of the particles which was stabilised by a subsequent cross-linking of BSA molecules with either BNBA and DMT-MM or genipin. Cross-linking by BNBA and DMT-MM served in this manuscript as the reference procedure since its whole procedure as its mechanism has formerly been described by LI *et al.* (Li *et al.* 2017). The MnCO_3_ core of the particles was dissolved by complexation of manganese(II) by EDTA resulting in Hb being surrounded by a shell of cross-linked BSA proteins. In order to dissolve MnCO_3_, the equilibrium between Mn^2+^ in solution and Mn^2+^ bound in solid MnCO_3_ was used for the synthesis procedure of the particles. The complexation of dissolved Mn^2+^ by EDTA must disturb this equilibrium. As a consequence of this disturbance, additional soluble Mn^2+^ is donated from solid MnCO_3_ in order to reestablish the equilibrium. Since EDTA is present in excess amounts, all soluble Mn^2+^ should be trapped thereby dissolving all solid MnCO_3_.

The determination of Hb concentrations in the stock solution, the first supernatant and the supernatants after each washing step showed that there was hardly any Hb traceable in the supernatants after each washing step. This result implied that Hb is tightly enclosed by MnCO_3_ and that possible pores inside of MnCO_3_ particles must be chiefly smaller than the diameter of Hb proteins. The resulting encapsulation efficiency of Hb inside MnCO_3_ particles varied from 39 – 82% depending on the amount of Hb initially added to the Na_2_CO_3_ and MnCl_2_ solutions. The size of the resulting particles did not differ in dependence on the amount of cross-linking reagent added before dissolution of the MnCO_3_ core. For genipin as cross-linking reagent concentrations of 1, 2, 3 and 4 mM were chosen, all leading to capsules measuring 3 – 4 μm in diameter as measured by DLS (s. Table 1).

**Table 1.**
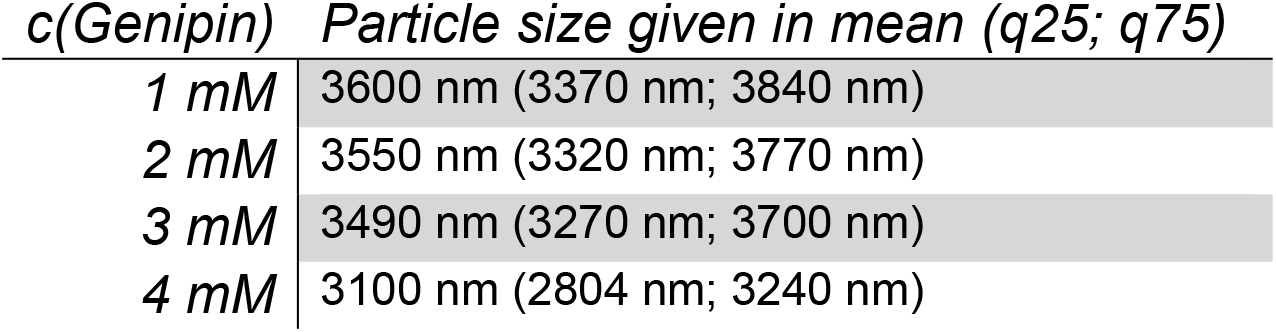
Particle sizes of HbOCs cross-linked by varying amounts of genipin as measured by DLS. Particle sizes are given as mean value and first (q25) and third (q75) quartile (n=3 each).

| *c*(Genipin) | Particle size given in mean (q25; q75) |
| --- | --- |
| 1 mM | 3600 nm (3370 nm; 3840 nm) |
| 2 mM | 3550 nm (3320 nm; 3770 nm) |
| 3 mM | 3490 nm (3270 nm; 3700 nm) |
| 4 mM | 3100 nm (2804 nm; 3240 nm) |

Cross-linking by BNBA and DMT-MM in DMSO also resulted in particles measuring approximately 3 μm in diameter assuming spherical shape of the microcapsules (Li *et al.* 2017), as seen in Figure 4. SEM images of BNBA-DMT-MM and genipin cross-linked particles were taken to depict the structure of the microparticles. Size measurement evaluated by SEM imaging yielded in particles that were seemingly one fourth to one seventh in size compared to measurements performed by DLS. Compared to a normal human RBC whose average size is about 8 μm in diameter and 2 μm in thickness, the prepared microcapsules have a volume of about 0.000113 fL, as measured by SEM, and RBCs are therefore approximately 800,000 times bigger with an average MCV of 94 fL (Diez-Silva *et al.* 2010). This small size hints at possible suitability for venoclysis.

**Figure 4.**
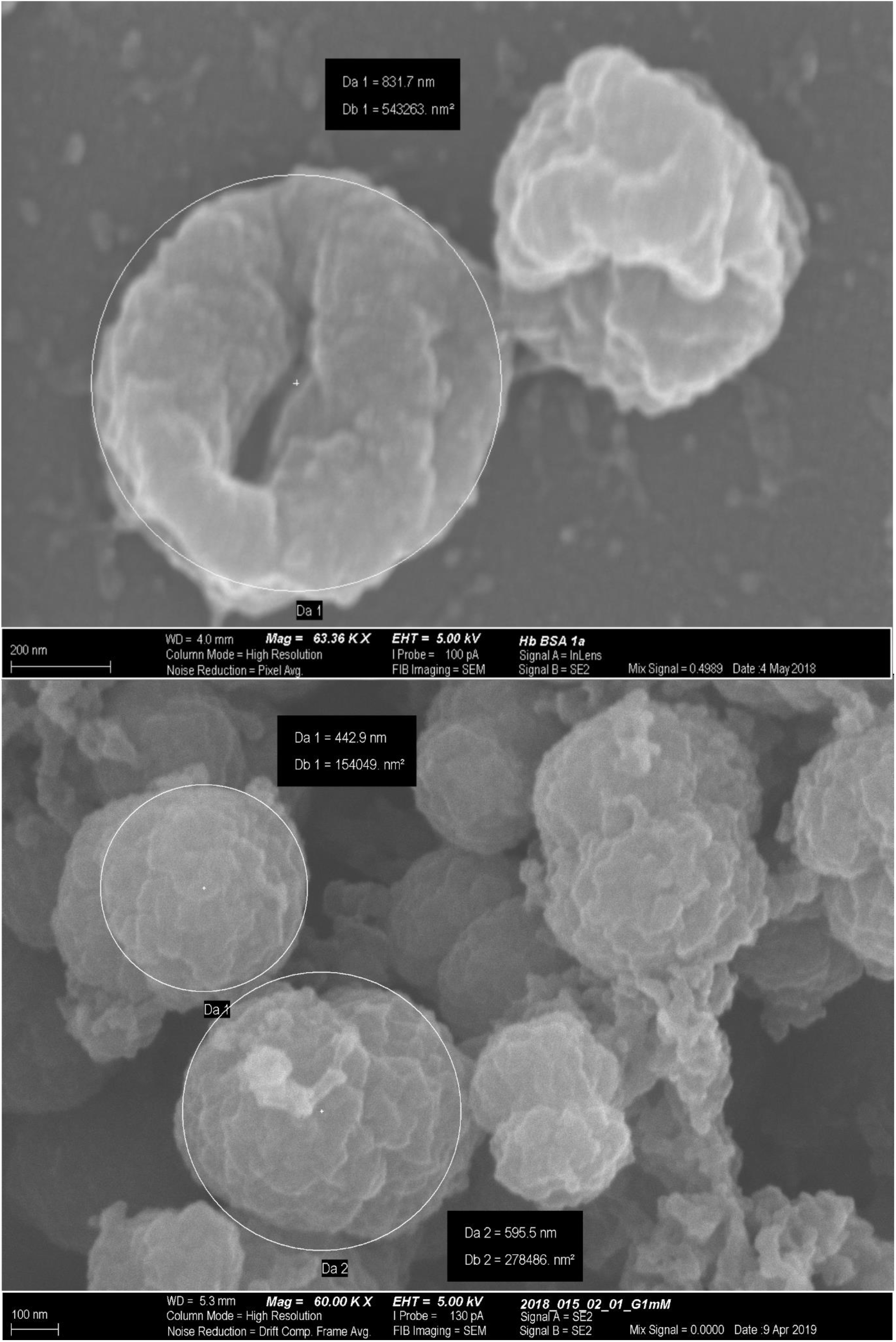
SEM images of HbOCs cross-linked by BNBA and DMT MM (top) and genipin (bottom). Disparity between DLS and SEM size measurements are due to aggregation of capsules at rest.

Oxygen binding capacity was measured by application of SDT as both reducing agent for methaemoglobin and de-oxygenation agent. The percentage of oxygenation was calculated from the total amount of Hb applied to the measuring chamber and the decrease in oxygen concentration. Our findings depict that both free Hb and encapsulated Hb were finally oxygenated up to 96%. (s. Figure 5).

**Figure 5.**
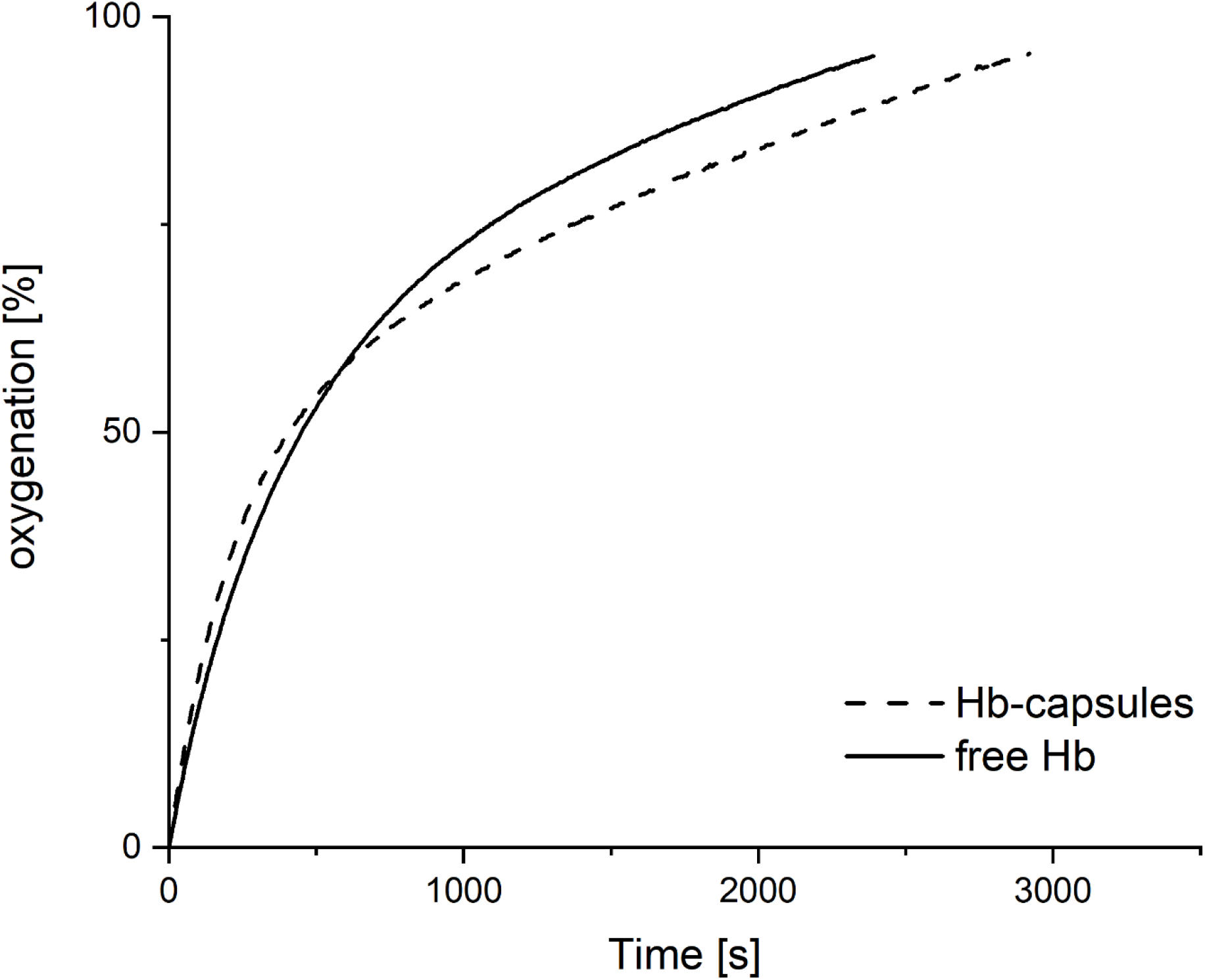
Determination of the oxygen binding capacity of HbOCs compared to the oxygen binding capacity of free Hb. By addition of SDT to HbOCs and Hb both were deoxygenated and reduced into the active ferrous forms before being injected into the chambers of the respirometer. For reasons of clarity and comprehensibility only one graph is shown exemplary.

In order to provide the purity of the samples, they were checked for possible noxious impurities resulting from the fabrication process. It was checked for residue Mn(II) and free Hb. First, the samples were washed three times with 30 mL Millipore water each. Then Na_2_CO_3_ was added to precipitate possible residue Mn(II). The supernatants were again checked for Hb by the SDS-Hb-method as described. Any tests failed in either finding Mn(II) or free Hb.

Measurements of LDH-activity before lysis showed slightly increased LDH-activities in supernatants taken from cells previously treated with HbOCs, where cells treated with 5 vol% of HbOCs released significantly (p = 0.014) less LDH than cells treated with 2.5 vol%, yet all LDH-activities were determined to be less than 15 U/L. However, after total lysis induced by Triton X-100, cells treated with 5 vol% of HbOCs displayed significantly (p = 4.2 ∙ 10^−8^) lower LDH-activities than any other group of cells tested (s. Figure 6). This was not the case when cells were incubated with an equal amount of free Hb (data not shown). Free Hb concentration was calculated as about 4.4 ∙ 10^11^ capsules per microliter (5 vol% HbOCs) based on SEM measurements (s. Figure 4). Comparison of relative cell damage revealed no damage of more than 5.2% in any treatment group (s. Figure 6), although treatment with 5 vol% HbOCs seems to have a minor but significant (p = 0.03) impact on relative cell damage. There is no evidence of any impact of the incubation duration on absolute cell damage. Remarkably, there is lower relative cell damage after three hours of incubation, although that group of cells already displayed lower total LDH-release. Nonetheless, this difference is still statistically insignificant. Compared to the ‘blank’ group, p is 0.26 for g-BSA, 0.60 for 2.5 vol% and 0.72 for 5 vol%. When g-BSA is used as reference p-values drop to 0.17 for 2.5 vol% and 0.03 for 5 vol%.

**Figure 6.**
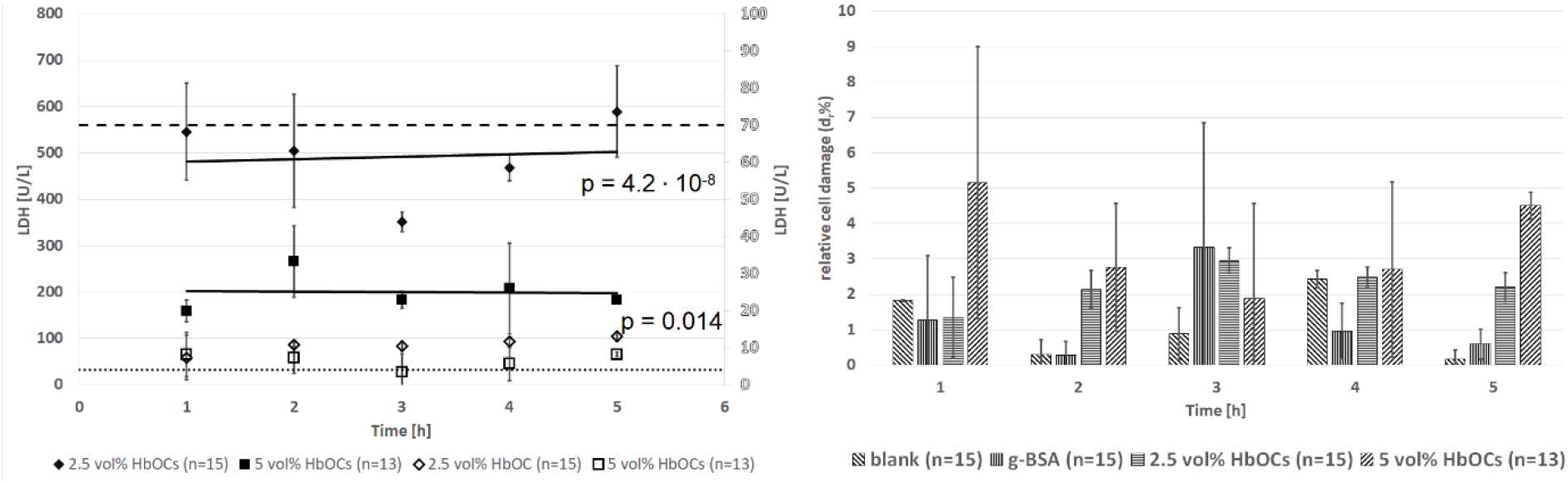
LDH-activities as a general marker of cell damage after treatment with BSA or HbOCs in different concentrations. A solution of 4.5 g/L glucose in Ringer’s solution served as blank. Either 3 mg/mL BSA (g-BSA) or a suspension of 2.5 vol% or 5 vol% HbOCs was added. Before (empty symbols; right y-axis) and after total lysis of cells (filled symbols; left y-axis) induced by addition of 200 μL 25% Triton X-100 as detergent. The dotted line represents the average LDH-activity of the untreated control cells (blank) before lysis. The dashed line represents the average LDH-activity of the untreated control cells (blank) after lysis (LDHmax). The relative damage has been calculated as described in methods. T-tests were performed between groups treated with 2.5 vol% and 5 vol% HbOCs. Data is shown +-1 standard deviation (5 vol% HbOCs n=13; all other n=15).

In order to investigate any interference of microcapsules with the LDH assay, standard testing solutions (DiaSys Diagnostic Systems GmbH, Holzheim, Germany) containing commonly tested parameters for clinical purposes with known and well defined activities or concentrations, respectively, were incubated 1:1 and 1:9 for 1 h at 37 °C with microcapsules suspension (5 vol%) or Ringer’s solution, respectively. (n = 3 each; s. Table 2 and Table 3). There were no significant differences in any parameters whether microcapsules had been added or not (p > 0.1 in any group, s. Table 2 and Table 3).

**Table 2.**
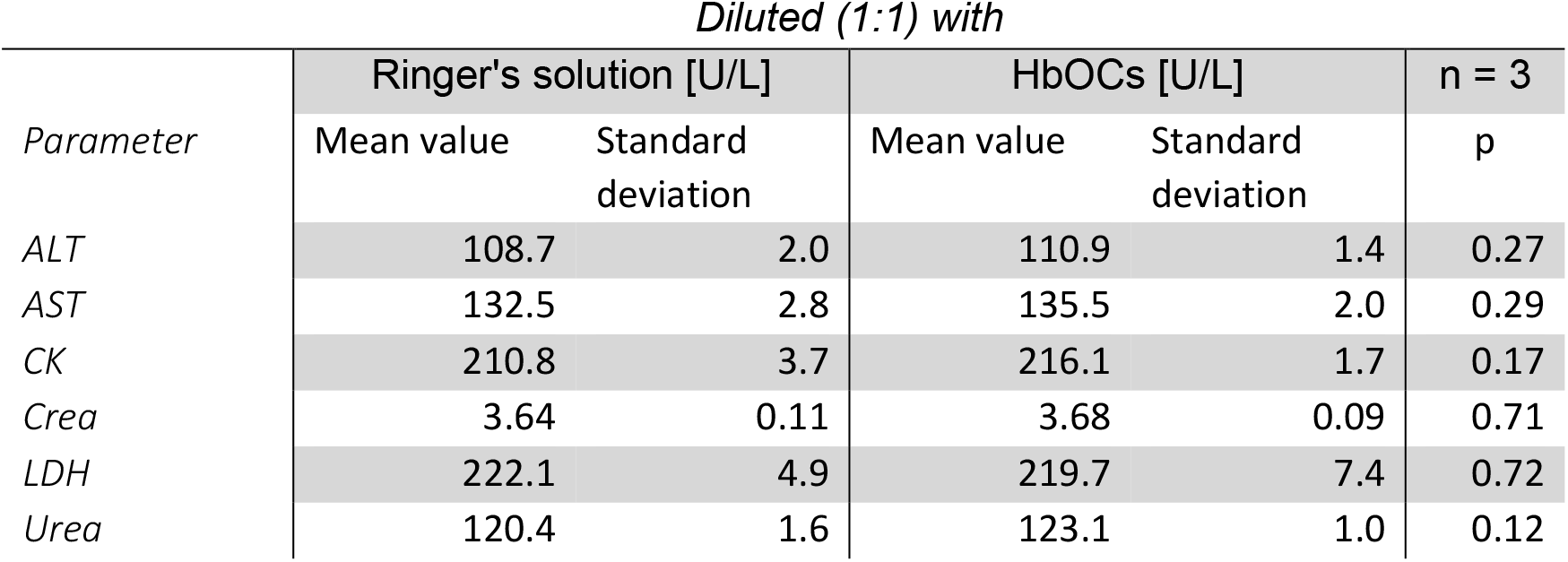
Testing on interaction between clinical standard parameters and prepared HbOCs. Ratio between standard testing serum and microcapsules suspension or Ringer’s solution, respectively, 1:1. All samples were incubated for 1 h at 37 °C (n=3).

**Table 3.** Testing on interaction between clinical standard parameters and prepared HbOCs. Ratio between standard testing serum and microcapsules suspension or Ringer’s solution, respectively 1:9. All samples were incubated for 1 h at 37 °C (n=3).

| Parameter | Diluted (1:9) with |  |  |  |  |
| --- | --- | --- | --- | --- | --- |
|  | Ringer's solution [U/L] |  | HbOCs [U/L] |  | n = 3 |
|  | Mean value | Standard deviation | Mean value | Standard deviation | p |
| ALT | 22.5 | 0.4 | 22.2 | 0.6 | 0.58 |
| AST | 26.8 | 1.3 | 27.3 | 0.4 | 0.62 |
| CK | 43.0 | 2.1 | 42.6 | 0.3 | 0.80 |
| Crea | 0.53 | 0.06 | 0.54 | 0.00 | 0.83 |
| LDH | 53.7 | 4.1 | 48.1 | 0.0 | 0.19 |
| Urea | 23.8 | 1.4 | 24.5 | 0.7 | 0.58 |

To investigate the difference of LDH release after incubation with 2.5 and 5 vol% of HbOCs, qNMR was performed for ruling out binding of Triton X-100 by HbOCs (s. Figure 7). Addition of 7.5 μL DMSO to 1 mL of 10% Triton X-100 in D2O results in 100 mM DMSO and 161 mM Triton X-100. After incubation of Triton X-100 solution with HbOCs at 37 °C no decrease in Triton X-100 concentration was determined. The resulting ratio between Triton X-100 and DMSO was determined to be 1.61 irrespectively of pre-incubation of Triton X-100 with capsules. These data clearly show, that microcapsules did not bind Triton X-100. Therefore, in the LDH assay (s. Figure 5), all cells were lysed with the same concentration of Triton X-100, irrespectively of the presence of no, 2.5 or 5 vol% of HbOCs. In conclusion, qNMR measurements of Triton X-100 did not further elucidate the difference in LDH release after treatment with different amounts of HbOCs, thus at the moment there is no valid explanation for this limitation of LDH-release, that is already observed within the first hour. Evaluation of interference of HbOCs and their delivered oxygen with (intra-)cellular signalling pathways could be the next step in understanding the limitation in total LDH-activities after treatment with higher doses of HbOCs.

**Figure 7.**
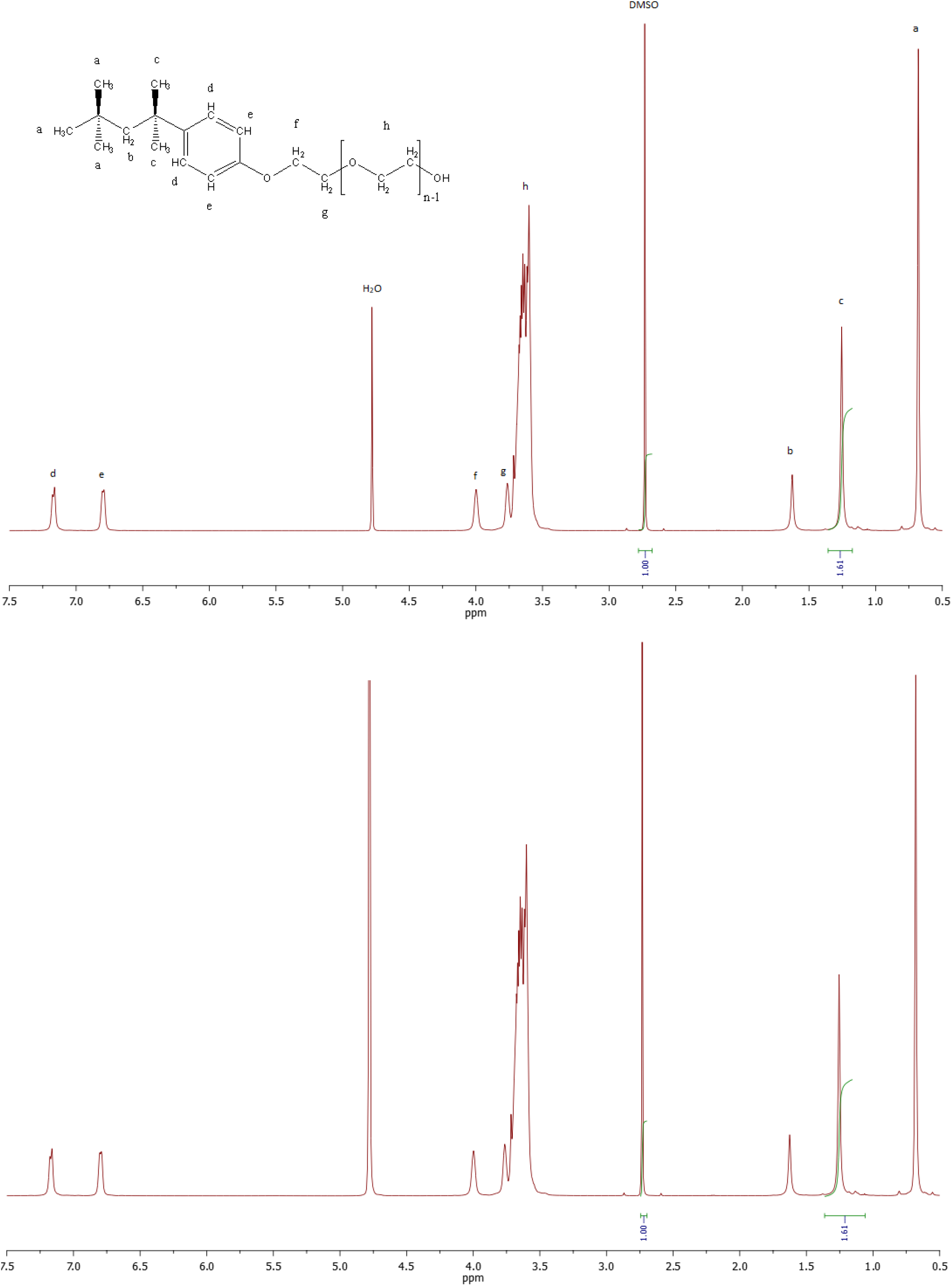
qNMR of 10% Triton X-100 with DMSO as internal standard. Without incubation (top) and after incubation over night with 5 vol% HbOCs (bottom). ^1^H-NMR (500 MHz, D_2_O) δ 7.17 (d, J = 7.2 Hz, 2H), 6.80 (d, J = 6.0 Hz, 2H), 4.00 (s, 2H), 3.76 (s, 2H), 3.73 – 3.43 (m, 36H), 1.63 (s, 2H), 1.26 (s, 6H), 0.68 (s, 9H).

Since BSA contains the basic amino acid lysin 59 times, it offers up to 60 amino-groups for cross-linking (Hirayama *et al.* 1990). Two amino groups are cross-linked by one molecule of genipin or other cross-linking agents, so that the maximum ratio of BSA:genipin equals 1:30. Using 1 mM genipin as described, equals a ratio of approximately 1:27 which is close to maximum cross-linkage. This results in an undesired linkage of two or more individual capsules.

In order to supress the artificial linkage of individual capsules, the applied amount of genipin was optimised. Further reduction of the amount of genipin used for cross-linking yielded in capsules with smaller diameters according to evaluation with DLS. A molecular ratio of 1:10 was found to result in two nearly Gaussian distributions with modes of 1230 nm and 2069 nm. Due to the small difference between the two modes, quartiles of the two distributions are overlapping. For the whole distribution only one percent of capsules is smaller than 690 nm and only one percent is bigger than 3400 nm. Further reduction of the BSA:genipin ratio down to 1:5 yielded in capsules displaying a similar distribution to those of ratio 1:10 but modes of the resulting fractions were torn apart to 989 nm and 2627 nm. On the one hand capsules were less cross-linked among each other, on the other hand the number of capsules linked incompletely also increased. Because of these findings, the amount of genipin cannot be further decreased. The optimal encapsulation efficiency was reached at concentrations of 0.1 M Na_2_CO_3_ and MnCl_2_ as well as 5 mg ∙ mL^−1^ Hb and 0.5 mg ∙ mL^−1^ BSA, yielding in 40 – 45% of encapsulated Hb. Further increase in Hb-concentration did not increase the absolute amount of encapsulated Hb. Further decrease in Hb-concentration led to higher encapsulation efficiencies but less encapsulated Hb in total.

Zeta-potential calculated for capsules prepared by the partly optimised method presented above yielded in ζ = −37.49 ± 1.25 mV given as mean ± standard deviation (n = 5). For the process of calculation, HbOCs were dissolved in 1 mL MilliQ water resulting in a concentration of approximately 1.2 vol% HbOCs. Each sample was calculated three times.

Albumin microcapsules containing haemoglobin prepared by the method presented can undergo oxidation of ferrous haemoglobin to ferric methaemoglobin just as it is naturally occurring in erythrocytes. It was shown that ferric methaemoglobin can be recovered easily by application of excess ascorbic acid or SDT in a non-enzymatic reaction, which is suggesting a long-term functionality of haemoglobin microcapsules prepared by the method presented (s. Figure 8). Due to the permeability of the albumin shell for small molecules, co-encapsulation of ascorbic acid together with haemoglobin is obviously not a mandatory prerequisite. Addition of ascorbic acid to a suspension of microcapsules right before application for venoclysis could present a suitable method to provide blood-levels of ascorbic acid high enough to re-reduce potentially oxidised ferric methaemoglobin.

**Figure 8.**
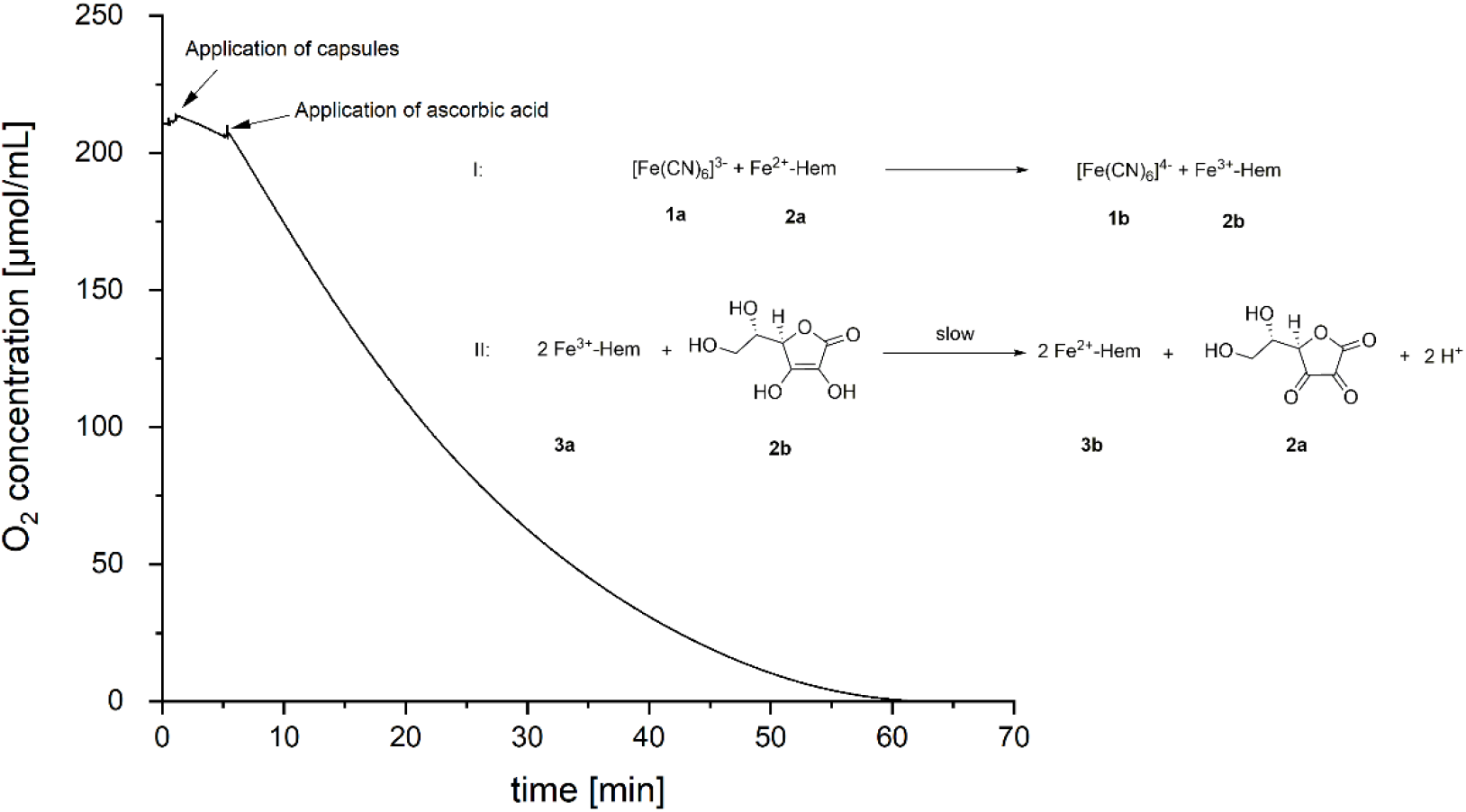
Testing on functionality by respirometry; Observation of change in oxygen concentration after application of HbOCs and ascorbic acid into the respirometer. I: Oxidation of ferrous haemoglobin (2a) by the ferricyanide anion (1a) to form the ferrocyanide ion (1b) and methaemoglobin (2b). II: Reduction of two equivalents of methaemoglobin iron (2b) by one molecule of ascorbic acid (3a) to form ferrous haemoglobin (2a) and dehydroascorbic acid (3b). For reasons of clarity and comprehensibility only one graph is shown exemplary.

## Conclusion

The preparation of Hb-microcapsules by the method presented leads to a promising type of artificial blood substitute. *In vitro* experiments showed a good biocompatibility of the applied microcapsules because any damage parameters remained to be almost silent. Nevertheless, the procedure is currently further optimised in order to receive particles with a somewhat smaller diameter. Therefore, interindividual cross-linking among capsules as well as aggregation of solitary capsules into larger clusters must be prevented for future use.

In any case, we demonstrated here for the first time, that the preparation of genipin-linked albumin microcapsules as potential artificial oxygen carriers is possible in absence of harmful cross-linking agents like glutaraldehyde.

## Acknowledgements

The authors thankfully express their gratitude to Mrs. Sylvia Voortmann of the Electron Microscopy Unit of the Imaging Center Essen (IMCES) for performance of SEM imaging, Mrs. Susanne Eitner from the Institute of Physiological Chemistry, University of Duisburg-Essen, University Hospital Essen (Essen) for performing Native PAGE and cell culture, Mrs. Birgit Podleska from the Institute of Physiological Chemistry, University of Duisburg-Essen, University Hospital Essen (Essen) for culturing cells and Mrs. Eva Hillen from the Institute of Physiological Chemistry, University of Duisburg-Essen, University Hospital Essen (Essen) for measurement of LDH-activity.

